# Isoform-resolved spatial transcriptomics on a lab-made high-density array via a single-chip NGS-TGS workflow

**DOI:** 10.64898/2026.03.15.711944

**Authors:** Zhiliang Yue, Min Liu, Yongqi Liu, Dongdong Lu, Menglong Zhang, Yuntong Wang, Yan Shi, Yuanyuan Miao, Suchen Wang, Yifan Jiang, Yue Wang, Jun Zhao, Naixu Liu, Chunyu Lv, Jixian Zhai, Bosheng Li

## Abstract

High-resolution spatial transcriptomics is still cost-prohibitive and dominated by short-read sequencing, limiting *in situ* detection of transcript structures. Here we present a low-cost, lab-made, high-density bead-in-microwell chip assembled by routine centrifuge and decoded by a tri-part combinatorial long-barcode strategy. The design provides 5.6×10^7^ barcode combinations across 4.1×10□ capture sites, reducing collisions while remaining tolerant to third-generation sequencing raw errors. Using a single-chip, dual-platform workflow, we split full-length cDNA for next-generation and third-generation sequencing within the same spatial coordinates. In an incompatible tomato-pepper graft, we identified interface-associated splicing reprogramming, including thousands of unannotated isoforms, enriched novel *CaGRP1* isoforms, and *SlPIP2* intron retention, supported by individual long-read sequences. In mouse embryos, we further demonstrate cross-species applicability by resolving unannotated *Col1a2* isoforms in oligodendrocyte progenitors and spatially restricted *Dalrd3* intron retention. This cost-efficient system broadens access to spatial full-length transcriptomics and may support diverse biological studies.

## Introduction

Life science research is undergoing a paradigm shift from tissue-level analysis to single-cell spatial resolution. The rise of spatial transcriptomics (ST) has enabled researchers to analyze broad-spectrum gene expression patterns while preserving original tissue context^1–6^. However, dominant high-resolution spatial technologies (e.g., 10x Genomics Visium, Slide-seq, and Stereo-seq) are largely built upon short-read next-generation sequencing (NGS) platforms^7–11^. This end-point counting approach, while excellent for cell clustering and differential expression analysis, overlooks the complexity of transcript structures. In eukaryotes, over 90% of multi-exon genes produce multiple isoforms through alternative splicing, alternative transcription start sites, or alternative polyadenylation, which often dictate protein functional diversity and cell fate^12–17^. Consequently, integrating spatial coordinates with isoform-resolved profiling has become a core frontier in spatial biology^6, 18–41^.

While the introduction of third-generation sequencing (TGS) offers a path toward spatial isoform analysis, integrating long-read sequencing (TGS) into high-density capture arrays faces severe challenges^42, 43^. First, there is a conflict between barcode capacity and chip area: achieving single-cell or subcellular resolution on millimeter-scale chips requires millions of capture sites. Currently, spatial barcodes, such as the 16-nt barcode used in 10x Visium, are short and susceptible to misassignment when sequencing errors occur. Second, the higher raw error rates of TGS platforms can interfere with the identification of short barcodes, causing some valid reads to fail spatial assignment. Furthermore, existing high-performance platforms often rely on expensive commercial chips or complex custom equipment, with high preparation costs per square millimeter, limiting their widespread adoption^44^. Consequently, current spatial transcriptomics research is often restricted to small sample sizes, making it difficult to exclude individual differences or technical noise through biological replicates^2^. Therefore, developing a low-cost and broadly deployable system that does not require specialized instrumentation is important for extending spatial omics from reference mapping to larger-scale biological and translational studies^45–47^.

Here, we present a high-density spatial transcriptomics system compatible with both NGS deep quantification and full-length sequence resolution. We introduced a tri-part combinatorial long barcode strategy based on self-assembled bead arrays. By generating over 5.6×10^7^ barcode combinations, this design supports approximately 4.1×10^6^ physical sites within a 6.8 mm × 6.8 mm capture area while maintaining a low simulated collision rate (6.98%) and improved tolerance to TGS read noise. Using this single-chip, dual-platform workflow, we constructed cross-species spatial full-length transcriptomic atlases, revealing spatial heterogeneity at the isoform level that is difficult to capture with traditional methods. In mouse embryos, we identified unannotated *Col1a2* isoforms in oligodendrocyte progenitor cells and detected *Dalrd3* intron retention at the single-molecule level. In the tomato-pepper incompatible graft model, we localized enriched novel *CaGRP1* isoforms and *SlPIP2* intron retention at the graft interface, suggesting spatially structured splicing reprogramming during tissue regeneration. In summary, we describe a lab-made, cost-efficient spatial transcriptomics platform for resolving transcript structures in complex microenvironments, with potential utility for studies of development, regeneration, and disease.

## Results

### A lab-made, high-density bead array enables isoform-resolved spatial transcriptomics via centrifugation-assisted assembly and tri-part barcoding

Commonly used spatial transcriptomics primarily relies on 150 bp paired-end NGS. Although this end-point counting strategy excels at gene quantification, its read-length limitations prevent the acquisition of full-length transcript information, leading to the loss of key structural variations such as alternative splicing, RNA editing, and fusion genes in the spatial dimension. To overcome this, we established a low-cost, lab-deployable, and TGS-compatible high-density spatial transcriptomics system and evaluated its manufacturability and performance in a standard laboratory setting. The core carrier of the system is a glass chip with a micro-processed surface etched with a microwell array. Each microwell has a diameter of 2.5 μm, a depth of 1.25 μm, and a center-to-center distance (pitch) of 3.5 μm. This design accommodates a single bead while providing mechanical constraints to prevent bead loss during fluidic operations. Within the 6.8 mm × 6.8 mm effective capture area, the array holds approximately 4.1×10□ physical capture points (spots, Fig. 1a), achieving high spatial resolution.

**Fig. 1.**
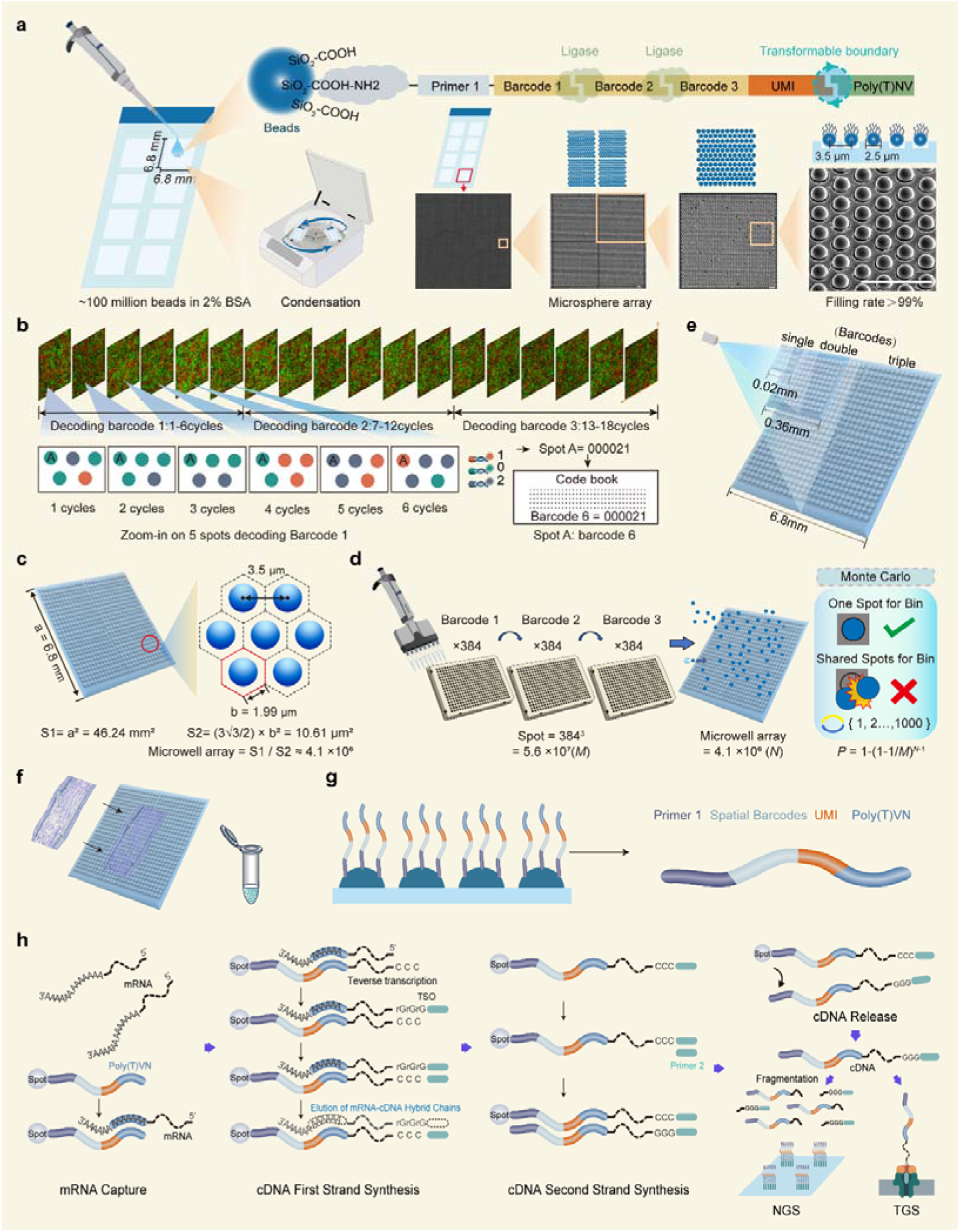
Construction and decoding of a full-length spatial transcriptomics platform. **a** Bead preparation and microwell array assembly. Silica beads are functionalized with sequencing primers, three-segment spatial barcodes, UMI, Poly(T)VN via split-pool ligation. The transformable boundary denotes a modular junction that enables substitution of the Poly(T)VN capture sequence with alternative capture moieties, including random primers or other custom-designed oligonucleotides. Beads are loaded onto chips containing ∼4.1×10□ microwells (2.5 μm diameter, 3.5 μm pitch) to achieve >99% occupancy. **b** HDST decoding workflow. Eighteen rounds of hybridization using Cy3/Cy5 channels resolve bead coordinates; each barcode segment is decoded in six rounds (3^6^ combinations) to distinguish 384 barcodes. **c** Schematic diagram of the calculation principle for the required spot counts in the microwell array. **d** Schematic diagram of the Monte Carlo calculation principle. **e** Schematic diagram of using single-, double-, and triple-part barcodes to cover chip area. **f** Tissue attachment and *in situ* cDNA synthesis following fixation and permeabilization. **g** Schematic of barcoded bead coating. **h** mRNA capture and library construction. Poly(T)VN captures mature mRNA for *in situ* reverse transcription, assigning spatial barcodes to cDNA for compatible NGS and TGS.

To overcome the dependence of traditional high-density chip preparation on expensive spotting equipment, we developed an efficient bead loading process based on centrifuge assisted assembly (detailed description in method). We first synthesized functionalized silica beads carrying sequencing primer Read 1, a tri-part spatial barcode, a molecular identifier (UMI), and a Poly(T)VN capture tail sequence. These beads were suspended in a 2% BSA solution and dropped onto the chip surface. After settling for 10 minutes, the chip was subjected to horizontal centrifugation using a standard benchtop centrifuge. The centrifugal force effectively overcame surface tension, actively pressing the beads into the depths of the microwells. After cleaning with nuclease-free water to remove excess beads, scanning electron microscopy (SEM) imaging showed a microwell filling rate exceeding 99% across the entire chip (Fig. 1a, Supplementary Fig. 1, Supplementary Video), with no obvious stacking or vacancies. This workflow simplifies chip loading through benchtop centrifugation and may facilitate implementation in standard laboratory settings.

For spatial barcode decoding, we employed a High-Density Spatial Transcript (HDST) strategy. Using decoding probes labeled with Cy3 and Cy5 fluorophores targeted to the barcode regions, we resolved the spatial coordinates of each bead *in situ* through 18 consecutive cycles of hybridization-imaging-stripping using a standard pathology scanner (e.g., 3DHISTECH Pannoramic MIDI, Fig. 1b and Supplementary Data 1). To ensure spatial address uniqueness at such high density and accommodate the high base error rates of TGS, we innovatively adopted a tri-part combinatorial barcode design. Within the 6.8 mm × 6.8 mm capture region of the chip, the center-to-center distance between adjacent spots in the matrix is 3.5 μm. Spatially, each spot is surrounded by six neighboring spots, and each physical capture site is shaped as a regular hexagon with an area of 10.61 μm². Across the full capture area of the chip (6.8 mm × 6.8 mm = 46.24 mm^2^), approximately 4.1×10□ physical capture sites can be accommodated (Fig. 1c). To ensure sufficient barcode uniqueness in this high-density array while also maintaining compatibility with the relatively high raw base error rate of TGS, we developed a tri-part combinatorial barcode strategy. Through three rounds of ligation in a 384-well plate, three sets of predefined unique oligonucleotide sequences (Barcodes 1-3) were combined, increasing the theoretical barcode diversity to 384³ = 5.6×10□ possible combinations (Fig. 1d). We then used a Monte Carlo simulation program to assess the uniqueness of this design (detailed description in method). When 5.6×10□ barcode combinations (*M*) were randomly assigned to 4.1×10□ capture sites (*N*), 1,000 simulation iterations indicated that the probability of barcode reuse (*P*) was only 6.98% (Fig. 1d). To further examine the value of the tri-part design, we compared the capture area supported by single-part and two-part barcodes while fixing the barcode reuse probability at 6.98% (*P* = 6.98%). The results showed that single-part (*M* = 384) and double-part (*M* = 384² = 147,456) designs could support only 27 and 10,676 capture sites, respectively, corresponding to effective chip areas of just 0.02 mm × 0.02 mm and 0.33 mm × 0.33 mm (Fig. 1e). These areas would likely be insufficient for the spatial coverage typically required by most biological tissue sections. Thus, the tri-part design substantially expands barcode capacity for large high-density fields, while its segmented structure may improve tolerance to long-read sequencing noise.

Finally, cryosections were attached directly to the prepared chip surface (Fig. 1f). After permeabilization, cellular mRNA was released and captured by the Poly(T)VN probes on the bead surface. Using template switching technology, full-length cDNA synthesis was completed *in situ* (Fig. 1g, h). During this process, the spatial barcode was permanently integrated into the 5’ end of the cDNA. The eluted full-length cDNA library was then split: one portion for NGS for high-depth gene quantification, and the other for TGS to directly read full-length transcript structures (Fig. 1h). This single-chip, dual-platform design unifies the gene expression atlas and the transcript isoform atlas within the same spatial coordinate system.

### Cross-species spatial atlases validate long-read compatibility and cross-platform label transfer

We subsequently applied this barcoding system to generate a high-resolution spatial transcriptomic atlas of an incompatible tomato-pepper graft (Supplementary Fig. 2) using both NGS and optimized TGS^48^ (Supplementary Data 2). Unsupervised clustering identified 17 transcriptionally distinct cell clusters, which were consolidated into 13 major cell types based on known marker genes. In the tomato section, seven cell types were annotated: epidermis, endodermis, vascular bundle, cambium, xylem, pith, and callus. In the pepper section, six cell types were identified: cortex, phloem, cambium, xylem, pith, and callus (Fig. 2a, Supplementary Fig. 3a and Supplementary Data 3). Following standardization and clustering of the NGS spatial data, we used cross-platform label transfer to map the TGS data onto the same reference, revealing broadly concordant cell-type distributions at the annotation level (Fig. 2b). Integration of both platforms using the Harmony algorithm demonstrated high comparability and successful fusion within a shared low-dimensional space (Fig. 2c).

**Fig. 2.**
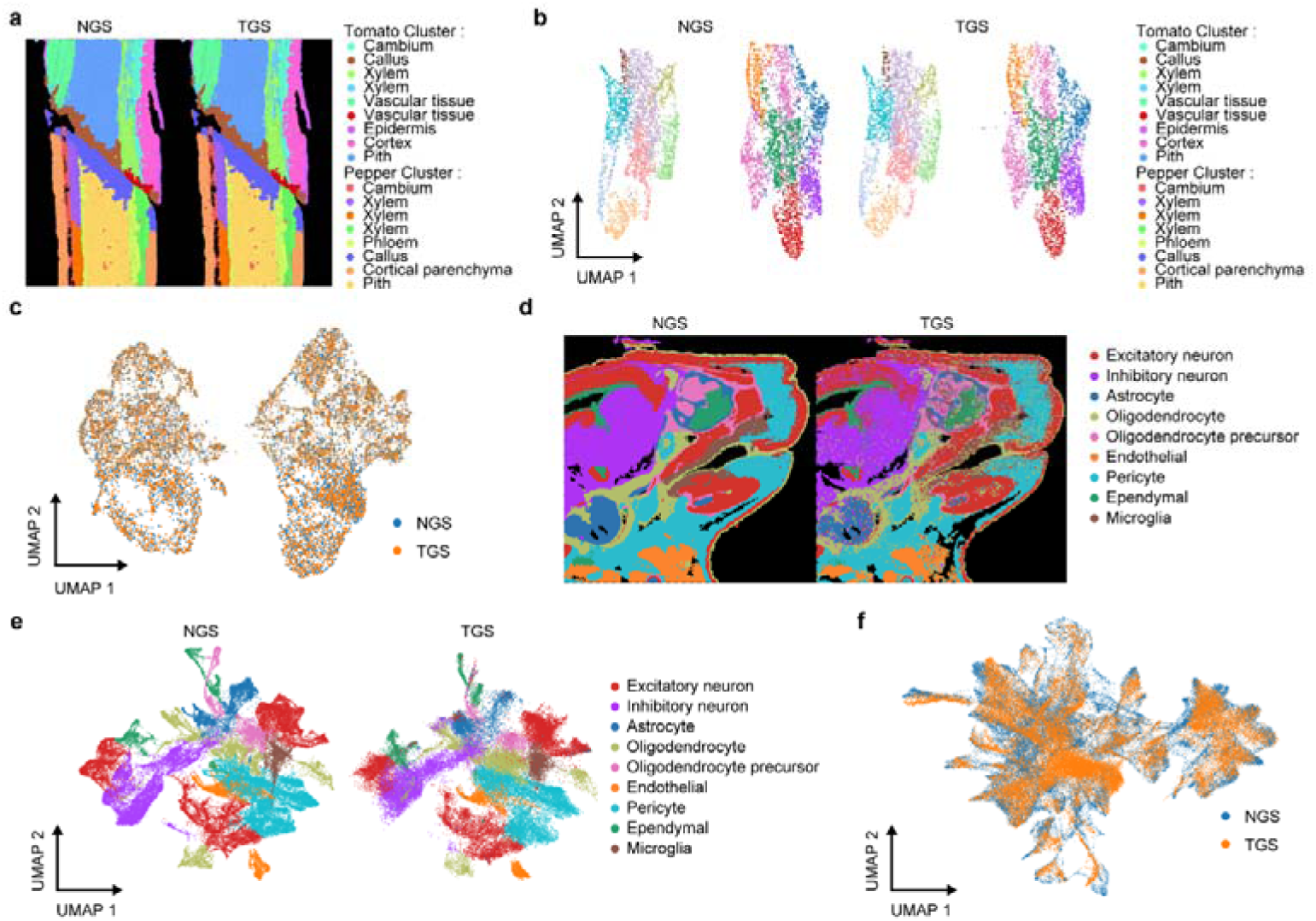
Spatial transcriptomic atlas generated by the full-length sequencing platform. **a** Spatial maps of tomato-pepper graft obtained via NGS and TGS, displaying identified cell types by artificial cell segmentation. Colors represent 17 spatial clusters corresponding to 13 tomato and pepper cell types. **b** UMAP projections of tomato-pepper graft data. NGS data were processed using standard pipelines, with cell types annotated by marker gene scoring. TGS data were mapped to the NGS reference space using tl.ingest for label transfer. **c** Integration of NGS and TGS tomato-pepper datasets in UMAP space following Harmony batch correction. **d** Spatial maps of mouse embryos obtained via NGS and TGS. Colors indicate 9 identified cell types. **e** UMAP projections of mouse embryo data. NGS data were normalized and clustered with cell type annotation performed via tl.score_genes. TGS data were mapped to the NGS reference using tl.ingest for cluster and label transfer. **f** UMAP projection of integrated mouse NGS and TGS data after Harmony batch correction demonstrating platform fusion.

We further extended this system to mouse embryos, performing dual-platform sequencing (Supplementary Data 2). In the mouse dataset, nine cell types were identified: excitatory neurons, inhibitory neurons, astrocytes, oligodendrocytes, oligodendrocyte progenitor cells (OPCs), microglia, endothelial cells, pericytes, and ependymal cells (Fig. 2d, Supplementary Fig. 3b and Supplementary Data 4). Similar to the plant model, cross-platform label transfer and Harmony-based integration verified consistent spatial distribution and reliable fusion between the NGS and TGS datasets (Fig. 2e, f). Together, these results support the cross-platform compatibility of our barcoding system and full-length library workflow. This analysis provides a practical framework in which NGS serves as a reference map for cell types and spatial structure, while TGS adds full-length transcript information within the same coordinates. Accordingly, our platform adds an isoform-resolved layer to standard spatial transcriptomic analysis while preserving the original tissue context.

### TGS resolves isoform-level spatial heterogeneity, revealing unannotated splicing and intron retention *in situ*

After obtaining reliable spatial cell-type maps and validating cross-platform comparability, we focused on a question that is traditionally difficult to answer with short-read spatial transcriptomics: whether the same gene exhibits differences in alternative splicing and transcript isoforms across different spatial microenvironments or cell types. Since TGS single-molecule reads can cover multiple splice sites and maintain long-range connectivity, we were able to incorporate transcript structure as part of the spatial molecular phenotype. This allowed for the *in situ* quantification and spatial projection of both annotated and unannotated isoforms, as well as typical splicing events such as intron retention (IR), within the same spatial coordinate system. Utilizing full-length transcript information, we compared transcripts with reference annotations and categorized them using the SQANTI3 methodology^49^. This classification included Full Splice Match (FSM) transcripts, where all splice junctions perfectly match the reference; Incomplete Splice Match (ISM) transcripts, which partially match consecutive junctions; and novel isoforms categorized as Novel In Catalog (NIC) or Novel Not in Catalog (NNC) based on the presence of known or novel donor and acceptor sites (Fig. 3a).

**Fig. 3.**
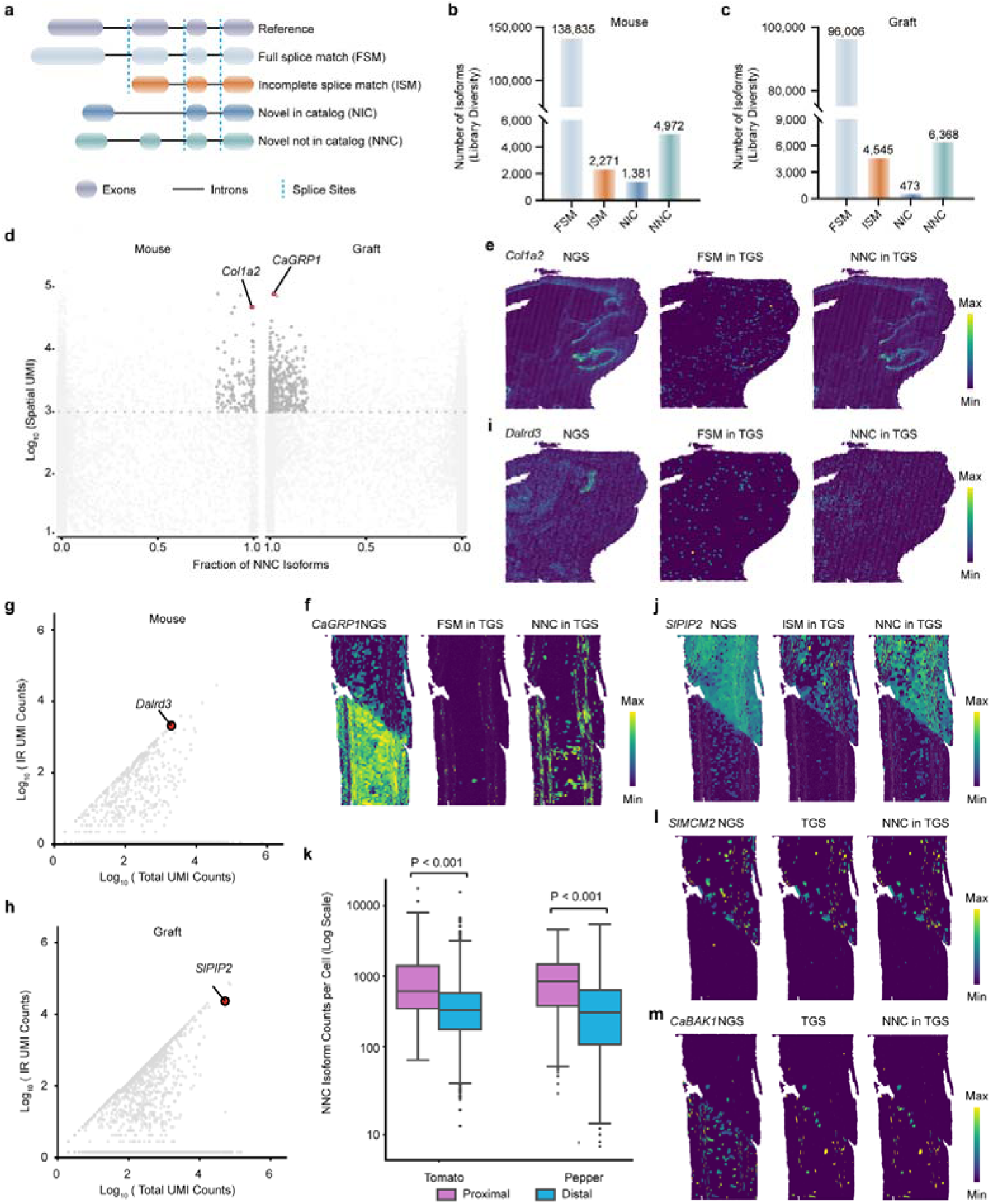
Spatial heterogeneity of transcript alternative splicing analyzed by TGS. **a** Schematic of transcript classification based on splice sites relative to reference annotations. Colored ellipses: exons; black lines: introns; blue dashed lines: alternative splicing sites. **b**, **c** Bar plots showing read counts for transcript categories in mouse embryo (**b**) and tomato-pepper graft (**c**) full-length datasets. **d** Scatter plots showing total gene expression versus the fraction of unannotated NNC (Novel Not in Catalog) isoforms for mouse (left) and tomato–pepper (right). Dark grey dots indicate top candidates filtered by a dual-threshold strategy (Total UMI >500–1000; NNC fraction >0.8). Highlighted dots identify *Col1a2* (mouse) and pepper *CaGRP1* (graft) as top-ranked genes. **e**, **f** Spatial projections of *Col1a2* (**e**) and pepper *CaGRP1* (**f**) isoforms across NGS and TGS platforms. **g**, **h** Identification of dominant IR targets. Scatter plots correlate total gene expression with IR isoform absolute abundance. *Dalrd3* (**g**) and tomato *SlPIP2* (**h**) are identified as high-confidence candidates (top-right quadrant), showing high transcriptional activity and significant unspliced transcript accumulation. **i**, **j** Spatial projections of *Dalrd3* (**i**) and *SlPIP2* (**j**) isoforms. **k** Box plot of NNC isoform counts per cell in proximal versus distal regions of the graft interface. **l**, **m** Spatial projections of interface-enriched genes *SlMCM2* (**l**) and *CaBAK1* (**m**) comparing NGS, TGS, and specific isoform distributions.

At the global transcript level, the number of detected transcript types varied between the mouse embryo (Fig. 3b and Supplementary Data 5) and tomato-pepper graft (Fig. 3c and Supplementary Data 5) datasets. FSM categories accounted for the vast majority of transcripts. Notably, 4,972 and 6,368 novel splicing forms not present in existing catalogs were detected in the mouse and graft datasets, respectively; the emergence of these unannotated forms suggests that transcript-level complexity at spatial resolution may be systematically underestimated. Crucially, our platform not only detects new isoforms but also links them to spatial enrichment patterns and cell-type contexts. Therefore, our approach may help prioritize recurrent transcript structures that are enriched in specific cell populations or spatial domains, over rare structures that may arise from low-abundance sampling noise. We further examined in the spatial dimension whether these transcripts are compatible with known tissue structures, developmental processes, or local microenvironmental features (Fig. 3b, c and Supplementary Data 5).

Subsequently, we statistically analyzed the proportion of unannotated NNC isoforms relative to total gene expression within the mouse and tomato-pepper graft spatial datasets. *Collagen type I alpha 2 chain* (*Col1a2*) in mouse and *Glycine-rich RNA-binding protein 1* (*CaGRP1*) in pepper were identified as the genes with the highest NNC proportions (Fig. 3d and Supplementary Data 6). These novel isoforms were validated by direct evidence from individual long-read sequences at single-molecule resolution (Supplementary Fig. 4a, b). These loci showed high expression levels while being composed predominantly of unannotated isoforms, suggesting incomplete reference annotation at these genes. The *Col1a2* gene encodes the α2 chain of type I collagen, a key component of the extracellular matrix (ECM)^50^. Observed spatially, unannotated NNC isoforms of mouse *Col1a2* were highly expressed in OPCs, consistent with the spatial pattern of NGS results. In contrast, the reference isoforms were detected at lower levels and lacked spatial expression specificity (Fig. 3e, Fig. S3a). The enrichment of *Col1a2* isoforms in OPCs is compatible with a role for *Col1a2* in the ECM microenvironment during oligodendrocyte lineage development^51^. In the tomato-pepper graft dataset, unannotated *CaGRP1* isoforms were enriched in vascular tissues, similar to reports linking related *GRP* genes to elongating tissues^52^. Conversely, reference isoforms were detected in smaller amounts and lacked spatial specificity (Fig. 3f, Supplementary Fig. 4b). This vascular enrichment raises the possibility that *CaGRP1*-related regulation is associated with elongating vascular tissues^52^.

To reduce the likelihood that apparent exon loss reflected RNA degradation rather than splicing, we screened for IR events at the gene level (Supplementary Data 7). We identified two genes with high accumulation ratios of unspliced intron transcripts: *DALR anticodon binding domain containing 3* (*Dalrd3*) in mouse embryos (Fig. 3g) and *PLASMA MEMBRANE INTRINSIC PROTEIN 2* (*SlPIP2*) in tomato (Fig. 3h). To validate the isoform analysis at the single-molecule level, we performed read-to-genome alignment visualization (using Minimap2) for mouse *Dalrd3* and tomato *SlPIP2*. While NGS showed high coverage across various exon regions, it was limited by read length and could not provide long-range connectivity information between exons, making it difficult to resolve complex combinations of isoforms. In contrast, TGS results provided individual reads spanning all splice sites from the 5’ end to the 3’ end, directly suggesting the full-exon collinearity of the transcript and demonstrating the advantages of single-molecule full-length sequencing (Supplementary Fig. 5).

In mouse *Dalrd3*, TGS captured full-length connectivity compatible with the coexistence of intron-retained and normally spliced transcripts (Supplementary Fig. 5a, red arrows). By contrast, the short-read NGS data did not preserve the same long-range connectivity, making these IR structures difficult to reconstruct directly (Supplementary Fig. 5a). In tomato, *SlPIP2* showed IR-related signals in both NGS and TGS datasets, indicating that both approaches can capture evidence of intron retention. However, while the NGS evidence is an assembly consensus from multiple sequences, the TGS advantage lies in its ability to sequence through the entire intron-retained form within a single read (Supplementary Fig. 5b). Furthermore, spatial mapping showed that TGS distinguished the spatial distributions of alternative splice forms, whereas NGS provided a more collapsed signal at the gene level (Fig. 3e, f, i, and j).

Utilizing the obtained full-length transcript information, we further analyzed the graft data containing two species. The callus and unconnected vascular bundles of the tomato and pepper components were defined as the interface-adjacent region (proximal), while the remaining cell types were defined as distal (Supplementary Fig. 6a). Differential gene analysis between the proximal and distal regions in the NGS results (Supplementary Data 8) revealed 443 upregulated genes in the tomato proximal region (log fold changes >1, *p* <0.05), primarily involved in cell cycle regulation (Supplementary Fig. 6b and Supplementary Data 9). In the pepper proximal region, 838 genes were upregulated, which were primarily involved in stress response (Supplementary Fig. 6c and Supplementary Data 10). Subsequently, analysis of isoform expression patterns revealed that unannotated novel isoforms were significantly enriched in the proximal regions of both species (Fig. 3k). To verify this, we performed spatial projections for the tomato proximal-enriched gene *SlMCM2* and the pepper proximal-enriched gene *CaBAK1*. Results showed that the NNC-type isoform expression pattern was consistent with the enrichment of all TGS isoforms in the proximal region (Fig. 3l, m, Supplementary Fig. 6d, e), suggesting that NNC-type isoforms account for a high proportion of TGS full-length transcripts and implying that the graft interface microenvironment is accompanied by considerable splicing reprogramming. Therefore, isoform-level spatial projection may help distinguish different transcript forms of the same gene across local cellular neighborhoods. This capability may be particularly useful for transcripts whose structural variation affects localization, stability, or putative function, and may provide an entry point for future splicing-spatial-phenotype studies.

## Discussion

The rapid development of spatial transcriptomics has been largely driven by commercial platforms; however, high implementation costs and closed technical systems often restrict the widespread adoption and flexibility of experimental designs. Here we describe an accessible spatial transcriptomics system based on a self-assembled bead array and centrifuge-assisted loading. Compared to 10x Genomics Visium (55 μm resolution) or technologies such as Slide-seq and Stereo-seq that require complex custom equipment or expensive instruments^7–11, 44^, our system achieves subcellular resolution using standard laboratory equipment and conventional reagents (2.5 μm beads, 4.1×10□ capture points). Furthermore, the split-pool bead synthesis strategy employed here imparts significant modular potential to the platform. While this study primarily demonstrates the capture of mRNA using Poly(T)VN, the transformable boundary incorporated into our ligation design allows for flexible replacement of capture sequences (Fig. 1a). This adaptable architecture could support future multi-omics extensions; for example, alternative capture chemistries may be incorporated for spatial proteomic or CRISPR-associated readouts.

Successfully implementing long-read spatial sequencing on ultra-high-density arrays requires precise barcode identification-a challenge exacerbated by the relatively high raw base error rates of TGS platforms. Our implementation of a tri-part combinatorial barcode strategy represents a key methodological innovation. Traditional spatial methods often rely on shorter, continuous barcodes that are prone to high collision rates or identification errors during TGS. By expanding the barcode space to over 5.6×10^7^ combinations, we reduced the simulated collision rate to 6.98% across a 6.8 mm × 6.8 mm field while increasing barcode capacity for long-read spatial analysis. By partitioning the barcode into multiple segments, this design is expected to improve error tolerance during demultiplexing in the presence of long-read sequencing noise. Together with the expanded barcode space and reduced barcode collisions, it may help mitigate the trade-off between spot density (spatial resolution) and decoding accuracy, providing a scalable framework for high-throughput TGS-based spatial analyses.

Beyond technical accessibility, this research addresses a critical blind spot in spatial biology: the inability of traditional short-read technologies to resolve complex transcript structures *in situ*. By integrating TGS with our high-density arrays, we revealed isoform-level heterogeneity that is difficult to resolve using NGS-based end-point counting alone. Our biological validations in both mammalian and plant models highlight the scientific value of this capability. In mouse embryos, we identified novel *Col1a2* isoforms specifically enriched in oligodendrocyte progenitor cells, suggesting that specific transcript variants may contribute to ECM remodeling during neurodevelopment. In the tomato-pepper incompatible graft model, we observed interface-associated splicing reprogramming, including enriched unannotated *CaGRP1* and *SlMCM2* isoforms and specific intron-retention events. These findings suggest that post-transcriptional regulation, such as alternative splicing, may contribute to localized responses during tissue regeneration and stress adaptation^46, 47, 53, 54^. The ability to map spatial landscapes at the full-length transcript level provides a useful framework for investigating how RNA structural variation relates to cell state and local microenvironments.

Although our dual-platform strategy successfully combines NGS’s depth with TGS’s structural resolution, it also reflects current limitations: the cost per data unit for TGS remains higher than for NGS, limiting the feasibility of relying solely on long-reads for high-depth spatial quantification. At present, our strategy utilizes TGS data as a structural layer superimposed on the quantitative layer provided by NGS. As TGS throughput increases and costs continue to decline, we anticipate a gradual shift in spatial transcriptomics toward a full-length sequencing paradigm. Additionally, while we have demonstrated the effectiveness of this system in frozen tissues, extending compatibility to Formalin-Fixed Paraffin-Embedded (FFPE) samples-the most abundant resource in clinical biobanks remains a priority for future optimization. In summary, by providing a low-cost and isoform-aware spatial transcriptomics solution, we hope to broaden access to spatial full-length transcript profiling for studies of development and disease^55, 56^.

## Methods

### Materials and Grafting Procedure

Tomato (*Solanum lycopersicum* cv. AC), pepper (*Capsicum annuum* L. CA59), and mouse (*Mus musculus*) were used in this study. For the plant models, three-week-old tomato and pepper plants were grafted at the second internode using a cleft-grafting method.

### Photolithographic Preparation of Microwell Arrays

High-resolution spatial capture arrays were fabricated on glass substrates using standard photolithography. Six-inch quartz wafers were ultrasonically cleaned in acetone, isopropanol, and deionized water (15 min each), followed by oxygen plasma treatment (100 W, 2 min). Negative photoresist (SU-8 2005, MicroChem) was spin-coated at 3000 rpm for 30 s to achieve a target thickness of 10 ± 1 μm. Wafers were soft-baked at 95 °C for 2 min. UV exposure (365 nm, 150 mJ/cm^2^) was performed using a chromium mask with a hexagonal array of 2.5 μm diameter holes and 3.5 μm center-to-center pitch. After a post-exposure bake (95 °C, 3 min), the wafers were developed in propylene glycol methyl ether acetate (PGMEA) for 1 min, rinsed with isopropanol, and nitrogen-dried. Stability was enhanced by a hard-bake at 150 °C for 30 min. To prevent non-specific bead adsorption, the chips were treated with Sigmacote (Sigma-Aldrich) for hydrophobic silanization. Finally, wafers were diced into standard 75 mm × 25 mm slides. These slides are commercialized by BMKMANU (catalog no. ST03020).

### Synthesis of Carboxylated Silica Microbeads

Uniformly sized carboxylated silica beads were synthesized via a sol-gel co-condensation reaction using a flow-focusing microfluidic device. The dispersed phase consisted of tetraethyl orthosilicate (TEOS), carboxyethylsilanetriol (CES), anhydrous ethanol, deionized water, and 0.01 M HCl, stirred for 3 h at room temperature for pre-hydrolysis. The continuous phase was HFE-7500 fluorinated oil with 2% (w/w) fluorosurfactant. The phases were converged in the microfluidic chip (dispersed: 150 μL/hr; continuous: 400 μL/hr) to generate stable droplets. Following incubation at 65°C overnight, the beads were demulsified with 20% (v/v) perfluorooctanol (PFO) and cleaned via gradient centrifugation (10,000 rpm, 5 min). Beads with a particle size of 2.5 μm were obtained via cell-mesh filtration. These beads are commercialized by BMKMANU (catalog no. ST03019).

### Bead Synthesis and Combinatorial Barcoding

Spatially barcoded beads were generated using a three-round split-and-pool combinatorial synthesis strategy. In the first round, 384 distinct 5’-amino-modified oligonucleotides (Sangon Biotech) were covalently coupled to carboxylated beads via carbodiimide-mediated condensation in 384-well plates. Each first-round oligo comprised an Illumina Read 1 sequencing primer (22 nt), a segment 1 barcode (17 nt), and a linker 1 sequence (5 nt). Following the initial coupling, beads were pooled, thoroughly mixed, and redistributed into a new 384-well plate for the second round of synthesis. In this round, segment 2 oligos (384 types), containing a linker 1 complementary sequence (5 nt), a segment 2 barcode (19 nt), and a linker 2 sequence (6 nt), were hybridized and enzymatically ligated to the bead-bound oligos. The split-and-pool process was repeated for the third round, where segment 3 oligos (384 types)-consisting of a linker 2 complementary sequence (6 nt), a segment 3 barcode (19 nt), a 12 nt Unique Molecular Identifier (UMI), and a 22 nt polyT(VN) tail-were ligated to extend the barcode diversity. The final theoretical complexity reached 384 × 384 × 384 and over 56 million unique spatial barcodes.

### Chip Assembly and Microbead Filling

Functionalized beads were resuspended in a high-density loading buffer (2% BSA) to ensure uniform distribution. The micro-well chips, fabricated via photolithography, were placed horizontally, and 100 μL of the bead suspension (∼50 million beads) was applied. Following a 10-minute sedimentation period, a horizontal centrifugal force of 1,000 × g was applied for 2 minutes using a swing-bucket rotor to drive the beads into the micro-wells. Post-centrifugation, the chips were washed three times with nuclease-free water to remove excess beads, achieving a filling rate exceeding 99% (Supplementary Video).

### *In situ* Decoding of Spatially Barcoded Beads

To resolve the spatial coordinates of the tripartite barcoded beads on the chip, we implemented a sequential hybridization-based decoding strategy utilizing a ternary encoding scheme. Each of the three barcode segments was represented by a 6-digit ternary code (e.g., ‘022101’), providing a potential code space of 3^6 = 729 unique combinations, which is sufficient to distinguish the 384 distinct sequences per segment. The decoding signals comprised two fluorophores (Cy3 and Cy5) and a null signal (blank), designated as ternary states 1, 2, and 0, respectively. For each segment, six rounds of hybridization were performed using specific pools of fluorescently labeled complementary probes (Pools 1-6 for Barcode 1, 7-12 for Barcode 2, and 13-18 for Barcode 3, Supplementary Data 1). Following each hybridization round, multi-channel fluorescence imaging was conducted via a high-resolution scanner to capture the signal distribution. The complete spatial decoding workflow encompassed 18 cycles of hybridization and imaging. An automated image processing pipeline was developed to reconstruct the barcode sequences (available at GitHub: https://github.com/lbs-lab/fluPictureProcess).

### Monte Carlo Simulation Program

To assess barcode uniqueness on high-density chips, we implemented a Monte Carlo simulation in Python (available at GitHub: https://github.com/lbs-lab/Monte-Carlo). The simulation performed 1,000 random-sampling iterations to model the random assignment of 5.6×10□ barcode combinations (*M*) to 4.1×10□ capture sites (*N*), and to estimate the probability (*P*) of at least one barcode duplication at a given number of sites. Based on this framework, we further compared the theoretical effective capture area of single-, double-, and triple-part barcode designs under a fixed duplication-probability threshold. In the simulation described above, the probability (*P*) of barcode duplication was estimated using the following equation:

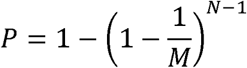

### NGS Short-read Data Processing and Expression Matrix Construction

Short-read data were processed using BSTMatrix (v2.4). The workflow included: fastq2BcUmi: parsing tri-part spatial barcodes and UMIs; Umi2Gene: aligning reads to reference genomes using STAR^57^ and counting at the gene level based on GTF annotations; LinkBcChip: mapping barcodes to physical coordinates using fluorescence decoding; and MatrixMake: performing UMI deduplication to output the final gene-by-barcode matrix and spatial coordinate files.

### TGS Long-read Data Processing

TGS data were processed using BSTMatrixONT (v1.4.2). The GenomeMap module utilized minimap2^58^ (map-ont mode) for alignment to the reference genome to generate sorted BAM files. The MatrixMake module then parsed barcodes and UMIs, integrated spatial information, and calculated effective UMI counts per barcode to generate the spatial expression matrix.

### Matrix Construction, Quality Control and Normalization

Count matrices were imported into a Scanpy environment, where each barcode was treated as a spatial sequencing unit (spot or cell). Spots with zero total counts were removed to ensure continuity. Library size normalization was performed using Scanpy^59^: scanpy.pp.normalize_total (target_sum = 1e4) followed by log 1 p transformation. Raw counts were stored for subsequent differential expression analysis.

### Spatial Mapping, Clustering Analysis and Tissue Annotation

NGS data were analyzed using Scanpy^59^. Highly variable genes (n = 3000) were identified, followed by PCA (n = 30) and neighborhood graph construction (n = 30). Clustering was performed using the Leiden algorithm (resolution = 0.8). Mouse embryo spots were annotated using marker genes (e.g., *Slc17a7*, *Gad1*) via tl.score_genes; graft data were annotated based on known markers reported in references.

### Cross-Platform Integration and Label Transfer

To ensure data consistency and eliminate platform-specific batch effects, we merged the TGS and NGS datasets and utilized Harmony for integration^60^, specifying the sequencing platform as a covariate. We further validated this integration by independently mapping the TGS data (as the query) onto the NGS reference UMAP embedding using scanpy.tl.ingest. This approach allowed for effective label transfer of cluster and cell type annotations, facilitating a precise cross-platform comparison.

### Isoform Detection, Classification, and Mapping

Transcript assembly and quantification for mouse TGS data were performed using IsoQuant^61^ (referencing GRCm38/Ensembl). SQANTI3^49^ was used for quality control and classification into FSM, ISM, NIC, and NNC categories.

### Integration of Isoform and Spatial Data

Read-isoform mapping from IsoQuant was combined with barcode spatial information to establish a spatial unit (spot or cell)-isoform-UMI association. UMI counts were aggregated by isoform and spatial unit using R scripts to generate 10x Genomics-style sparse matrices (.mtx) for spatial isoform usage analysis.

### Global Assessment of Cross-Species Intron Retention Events

A statistical pipeline in R was developed to quantify IR distribution. Isoform abundance matrices were loaded via Seurat^62^ and associated with SQANTI3 results. For each gene, Total Counts and IR-specific Counts were calculated.

### Global Visualization and Statistical Analysis

Cross-species IR abundance profiles were constructed using ggplot2. Total and IR expression levels were transformed and displayed as scatter plots, highlighting *Dalrd3* (mouse) and *SlPIP2* (tomato). The patchwork package and geom_text_repel were used for multi-species panel integration and gene annotation.

### Isoform-Level Spatial Projection and Expression Analysis

Targeted genes were analyzed by structural category. Custom Python scripts integrated isoform UMI counts into AnnData objects. Spatial expression preferences were visualized using scanpy.pl.spatial, projecting total gene expression and FSM/NNC isoform distributions simultaneously.

### Structural Validation via IGV

Key isoforms were validated using Integrated Genome Viewer^63^. Sorted BAM files and reference GTFs were loaded into separate tracks to illustrate long-read spanning of complex exon structures. High-resolution images were exported as evidence of isoform identification reliability.

## Supporting information

Supplementary Tables, and will be used for the link to the file on the preprint site.

Supplementary Video, and will be used for the link to the file on the preprint site.

## Data availability

The raw sequencing data generated during this study including mouse and plant grafting are available in the National Center for Biotechnology Information (NCBI) Sequence Read Archive (SRA) under BioProject accession number PRJNA1355050. The processed ST datasets, including gene expression matrices and isoform quantifications, have been deposited in the NCBI Gene Expression Omnibus (GEO) under accession number GSE316789 (token: wlyryikopnunzkp).

## Code Availability

The Monte Carlo simulation code for barcode-collision and spatial-capacity evaluation is available on GitHub (https://github.com/lbs-lab/Monte-Carlo). The automated image-processing pipeline for barcode reconstruction is also available on GitHub (https://github.com/lbs-lab/fluPictureProcess).

## Acknowledgements

We thank the Single-Cell and Single-Molecule Core Facility, Institute of Advanced Agricultural Sciences, Peking University, for providing the spatial transcriptomics and third-generation sequencing data. This work was supported by Shandong Provincial Natural Science Foundation (grant nos. SYS202206, ZR2023QC026 and ZR2023QC106), the National Key Research and Development Program of China (grant SQ2025YFE0201846), the National Natural Science Foundation of China (grant nos. 32200249 and 32170574), the Key Research and Development Project of Shandong Province (Agricultural Variety Improvement Project of Shandong Province, 2025LZGC004), and the Taishan Scholars Program and Yuandu Scholars Program.

## Author Contributions

Author contributions: B.L., J.Z., Z.Y., M.L. and Y.L. conceived of the project and designed the experiments or bioinformatics analysis; Z.Y., M.Z., Y.S., Y.M., Y.J and Y.W. conducted the majority of experiments, S.W. provided next-generation sequencing data, D.L. provided third-generation sequencing data, Y.L. and Y.W. performed the bioinformatics analysis; N.L drew the schematic diagrams; J.Z. worked for the SEM; C.L. prepared the plant materials; B.L., Z.Y. and Y.L. wrote the manuscript with input from all co-authors.

---

**Supplementary Fig. 1.**
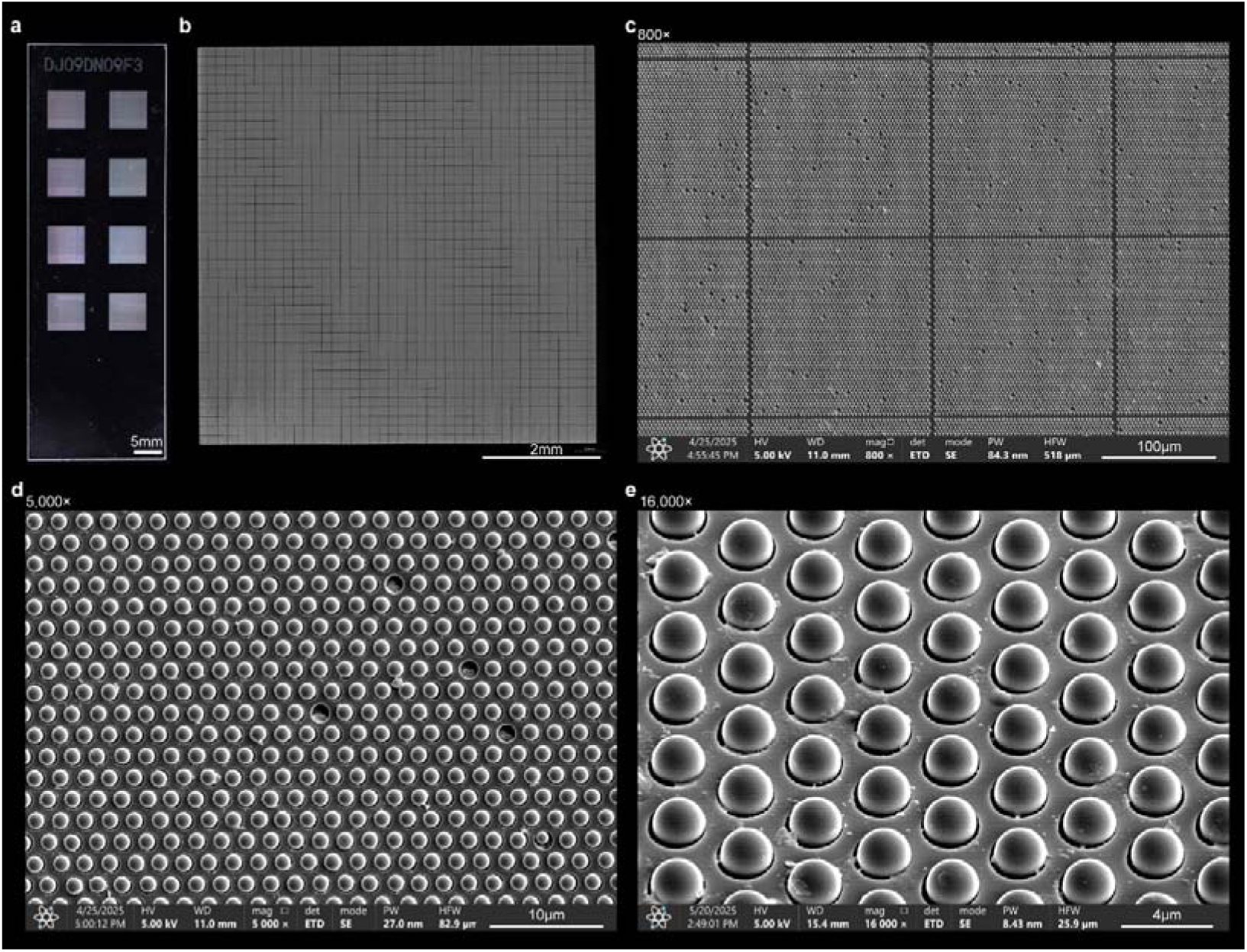
Structural characterization and hierarchical visualization of the chip. **a** Macroscopic photograph of the fabricated chip, showcasing the overall architecture. **b** Representative microscopic image of the effective capture area (6.8 mm × 6.8 mm). **c-e,** Scanning electron microscopy (SEM) images of the chip surface at increasing magnifications demonstrate high-uniformity loading process, with the microwell filling rate exceeding 99% across the entire chip area. Scale bars are provided in the bottom.

**Supplementary Fig. 2.**
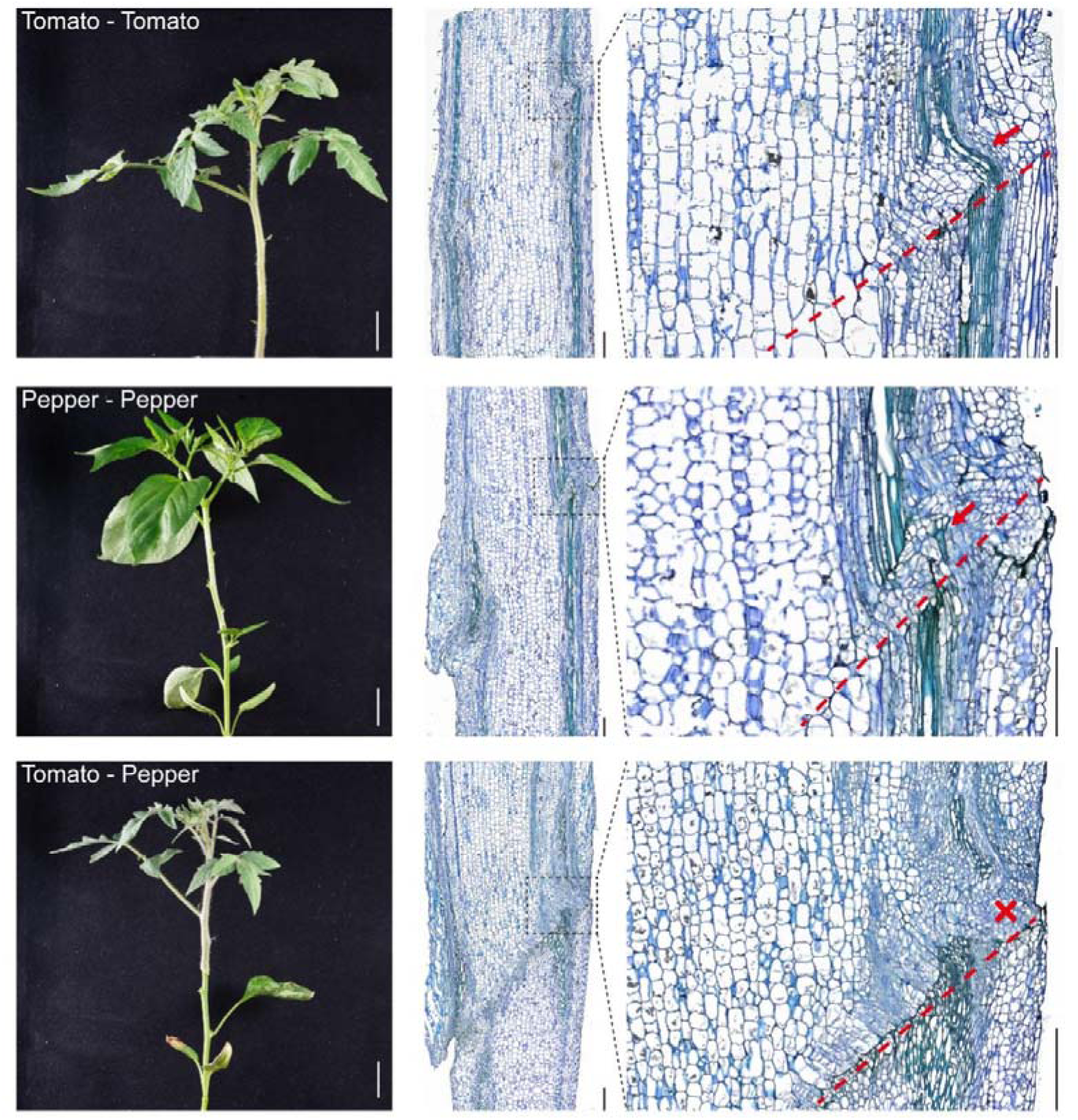
Incompatibility of tomato-pepper grafts. **Left**, phenotypic observations of tomato self-grafts, pepper self-grafts, and tomato-pepper heterografts at 10 days after grafting (DAG). Scale bar, 3 cm. **Middle**, Toluidine Blue O (TBO) staining of paraffin-embedded longitudinal sections at the graft interface; the presence of vascular bundles on both sides indicates mid-plane sectioning. **Right**, magnified views of the regions boxed in black dashed lines. Successful vascular reconnection (red arrows) between scion and rootstock is observed in self-grafts at the interface (red dashed line), while tomato-pepper heterografts exhibit a failure of vascular connection (red crosses). Scale bars, 100 μm.

**Supplementary Fig. 3.**
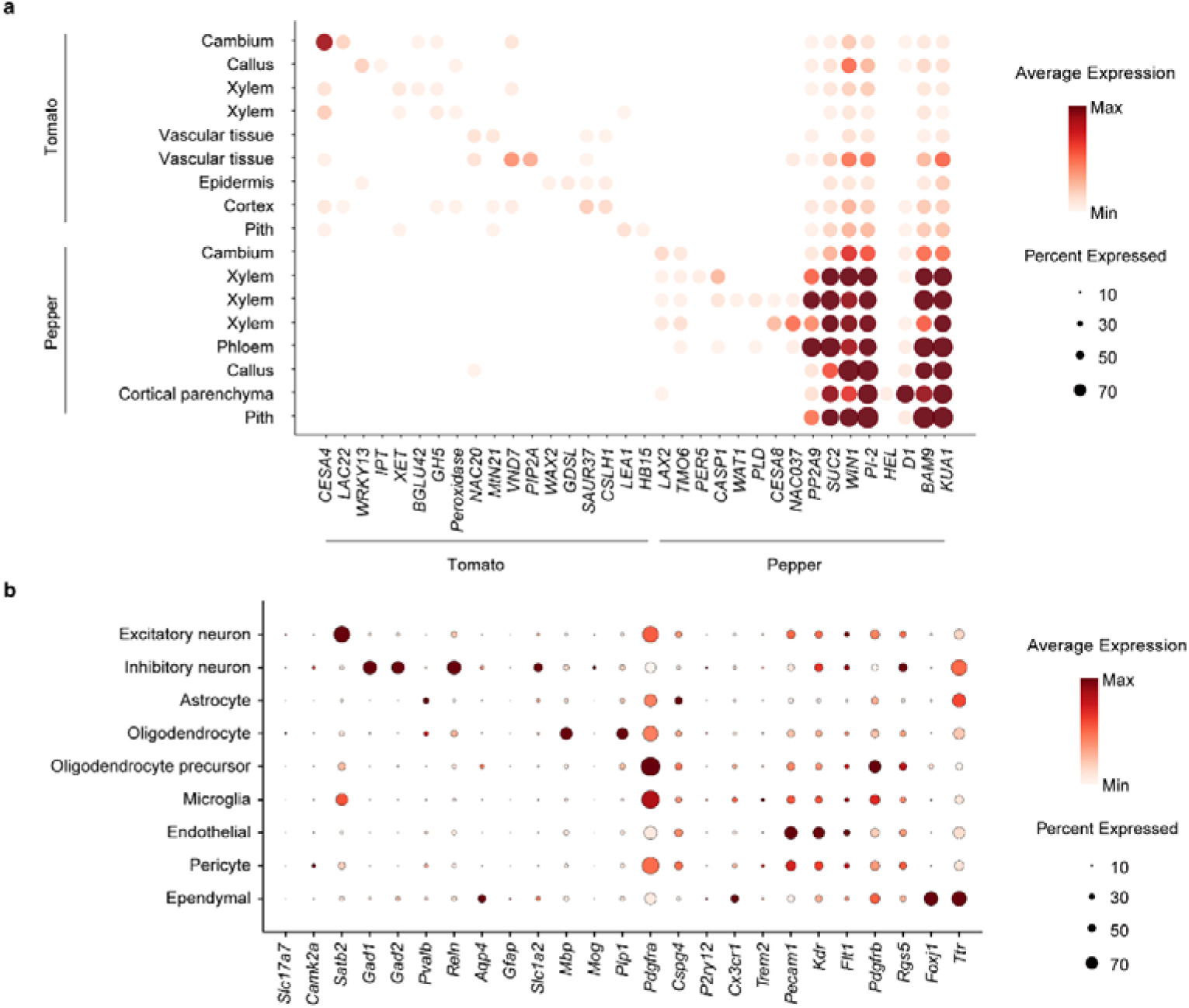
Bubble plots for the cell type-specific marker genes, with bubble size corresponding to gene expression levels in each cluster, for **a**, Tomato-Pepper graft, **b**, Mouse embryo spatial transcriptomic maps.

**Supplementary Fig. 4.**
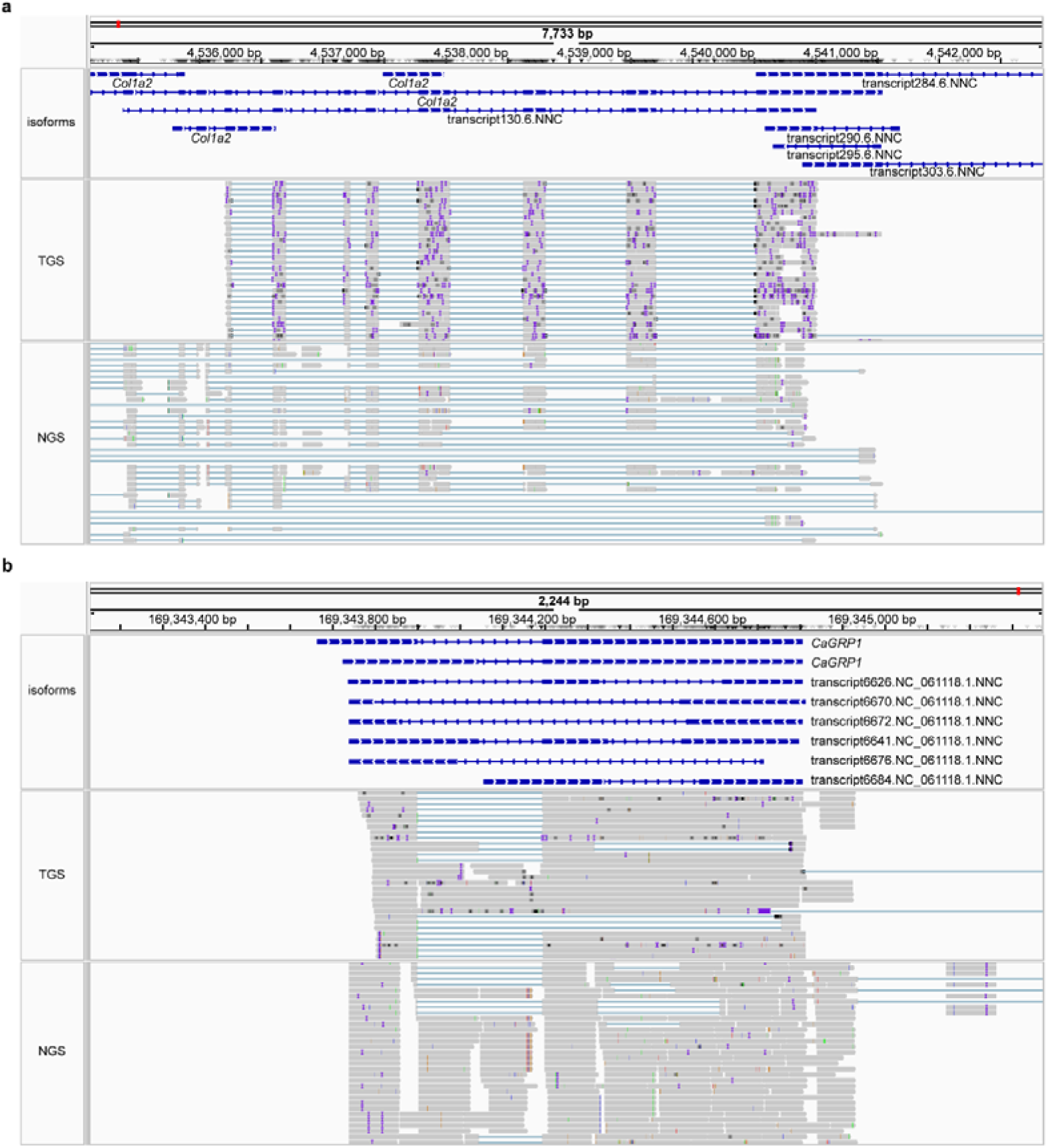
IGV-based validation of NNC via NGS and TGS. **a**, **b** Integrative Genomics Viewer (IGV) tracks comparing NGS and TGS detection of NNC isoforms for mouse *Col1a2* (**a**) and pepper *CaGRP1* (**b**). Top panels display schematic isoform structures with exons (blue bars) and introns (blue lines). Middle panels show individual TGS long-reads, where each row represents a single read. Bottom panels show NGS short-reads (gray bars with white gaps indicating gaps between paired-end or split reads) aligned to the reference transcripts. Isoform classifications are indicated following the transcript identifiers.

**Supplementary Fig. 5.**
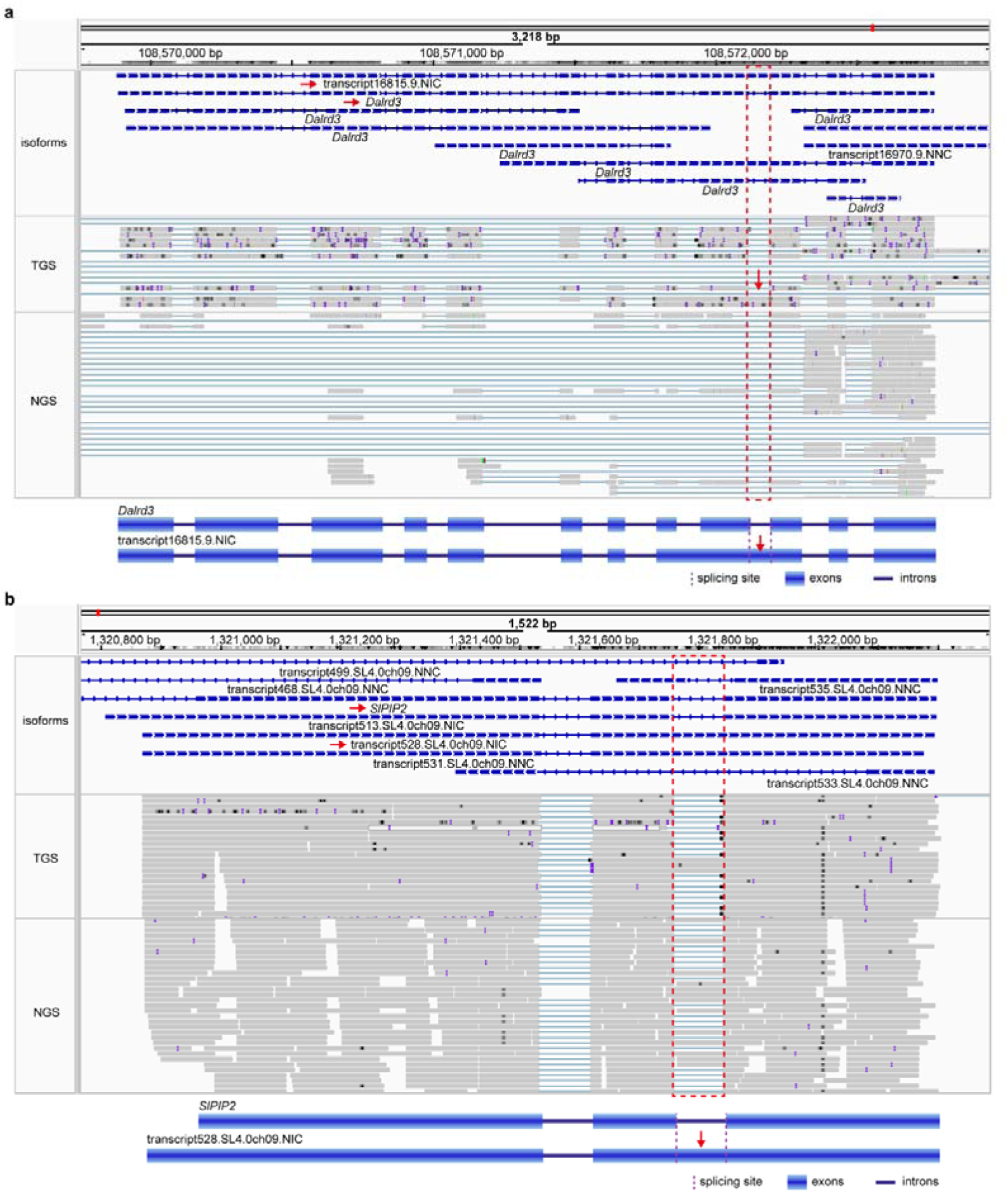
IGV-based validation of IR via NGS and TGS. **a**, **b** IGV tracks comparing NGS and TGS detection of IR isoforms for mouse *Dalrd3* (**a**) and tomato *SlPIP2* (**b**). Top panels show isoform schematics (blue bars, exons; blue lines, introns). Middle panels display individual TGS long-reads. Bottom panels show fragmented NGS reads aligned to reference transcripts. Red dashed lines highlight the retained regions where TGS and NGS coverage differ. Structural schematics of the IR isoforms are provided below the IGV plots.

**Supplementary Fig. 6.**
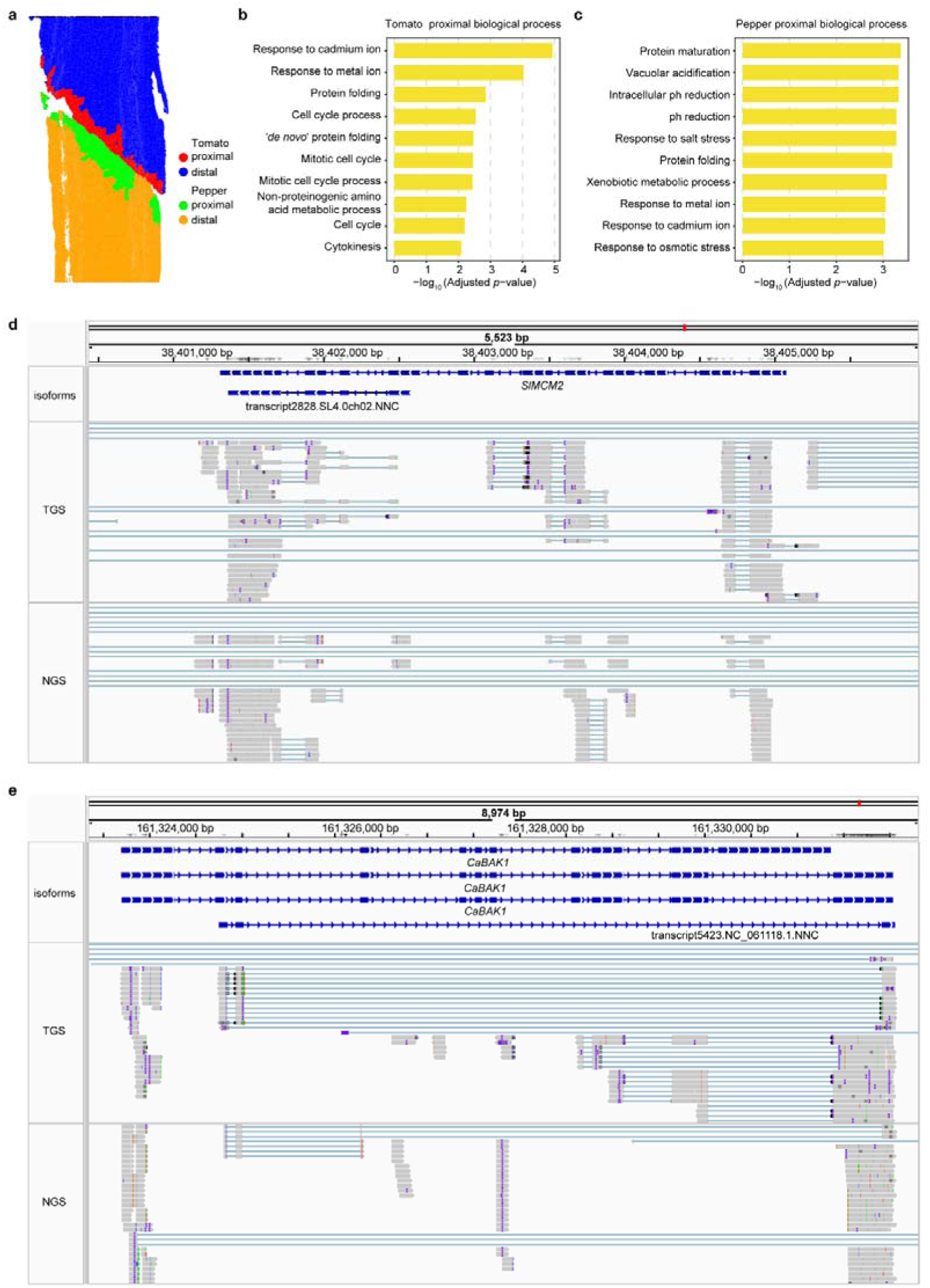
Spatial heterogeneity and functional analysis of transcripts at the graft interface. **a** Schematic of cell type extraction from proximal and distal regions of the tomato-pepper graft interface. **b**, **c** GO enrichment analysis of genes enriched in the proximal tomato (**b**) and pepper (**c**) interface regions. **d**, **e** IGV tracks comparing NGS and TGS reads for interface-enriched genes *SlMCM2* (tomato, **d**) and *CaBAK1* (pepper, **e**). Top panels show isoform schematics (blue bars, exons; blue lines, introns). Middle panels display individual TGS long-reads, while bottom panels show fragmented NGS reads with white gaps representing discontinuous alignment. Red dashed lines highlight differential exon retention regions captured by TGS versus NGS.

